# Late Anisian microbe-metazoan build-ups (“stromatolites”) in the Germanic Basin – aftermath of the Permian – Triassic Crisis

**DOI:** 10.1101/2021.03.15.435468

**Authors:** Yu Pei, Jan-Peter Duda, Jan Schönig, Cui Luo, Joachim Reitner

## Abstract

The so-called Permian – Triassic mass extinction was followed by a prolonged period of ecological recovery that lasted until the Middle Triassic. Triassic stromatolites from the Germanic Basin seem to be an important part of the puzzle, but have barely been investigated so far. Here we analyzed late Anisian (upper Middle Muschelkalk) stromatolites from across the Germanic Basin by combining petrographic approaches (optical microscopy, micro X-ray fluorescence, Raman imaging) and geochemical analyses (sedimentary hydrocarbons, stable carbon and oxygen isotopes). Paleontological and sedimentological evidence, such as *Placunopsis* bivalves, intraclasts and disrupted laminated fabrics, indicate that the stromatolites formed in subtidal, shallow marine settings. This interpretation is consistent with δ^13^C_carb_ of about −2.1‰ to −0.4‰. Occurrences of calcite pseudomorphs after gypsum suggest slightly evaporitic environments, which is well in line with the relative rarity of fossils in the host strata. Remarkably, the stromatolites are composed of microbes (perhaps cyanobacteria and sulfate reducing bacteria) and metazoans such as non-spicular demosponges, *Placunopsis* bivalves, and/or *Spirobis-like* worm tubes. Therefore, these “stromatolites” should more correctly be referred to as microbe-metazoan build-ups. They are characterized by diverse lamination types, including planar, wavy, domal and conical ones. Microbial mats likely played an important role in forming the planar and wavy laminations. Domal and conical laminations commonly show clotted to peloidal features and mesh-like fabrics, attributed to fossilized non-spicular demosponges. Our observations not only point up that non-spicular demosponges are easily overlooked and might be mistakenly interpreted as stromatolites, but also demonstrate that microbe-metazoan build-ups were widespread in the Germanic Basin during Early to Middle Triassic times. In the light of our findings, it appears plausible that the involved organisms benefited from elevated salinities. Another (not necessarily contradictory) possibility is that the mutualistic relationship between microbes and non-spicular demosponges enabled these organisms to fill ecological niches cleared by the Permian – Triassic Crisis. If that is to be the case, it means that such microbe-metazoan associations maintained their advantage until the Middle Triassic.

## Introduction

Microbes have dominated the biosphere for most of Earth’s history. This is for instance impressively documented by the widespread occurrence of microbialites across the breadth of time, with the oldest unambiguous records dating as far back as ca. 3.5 billion years ago (Lowe 1980; Walter et al. 1980; Grotzinger & Knoll 1999; Riding 2000; Arp et al. 2001; Duda et al. 2016; Mißbach et al. 2021). Hence, microbialites are important archives and provide valuable insights into the co-evolution of life and Earth through geological time because of their large spatial and temporal extent.

Principal categories of microbialites include stromatolites, thrombolites, dendrolites and leiolites (Kalkowsky 1908; Aitken 1967; Burne & Moore 1987; Riding 1991; Braga et al. 1995). Among these, stromatolites are likely the most famous type. The name is actually the translation of the German term “Stromatolith”, which was coined by Kalkowsky back in 1908. Kalkowsky introduced the term to describe layered carbonate structures in the Lower Triassic Buntsandstein of central Germany, and remarkably already proposed a biological origin of these fabrics. Following work showed that various biological processes are involved in stromatolite formation, including microbial mat-related mineral precipitation and/or trapping and binding of detrital sediments (Awramik et al. 1976; Reitner 2011; Riding 2011; Suarez-Gonzalez et al. 2019).

Ancient stromatolites from peritidal to shallow subtidal settings are commonly supposed to represent cyanobacterially dominated biofilms and microbial mats (see Monty 1977; Arp et al. 2001), analogous to modern stromatolites (Walter 1972; Dravis 1983; Arp et al. 1999; Dupraz et al. 2013). However, stromatolites can also be formed by light-independent chemolithotrophic or heterotrophic microbial communities, as for instance observed in caves and/or deep-water environments (Playford et al. 1976; Cox et al. 1989; Böhm & Brachert 1993; Heim et al. 2015, 2017). Particularly noteworthy are stromatolites formed by syntrophic consortia of anaerobic methane-oxidizing archaea and sulfate reducing bacteria (Reitner et al. 2005a, b; Arp et al. 2008). Notably, stromatolites do not have to be exclusively microbial in origin but can also consist of microbial mats and metazoans, e.g. sponges (Reitner 1993, Reitner et al. 1995; Riding & Zhuravlev 1995; Szulc 1997; Bachmann 2002; Luo & Reitner 2014, 2016; Lee & Riding 2020; Kershaw et al. 2021). Such complex structures, defined as microbe-metazoan build-ups, are easily overlooked due to the hardly identifiable morphological characters of non-spicular (“keratose”, see below) demosponges involved (Luo & Reitner 2014, 2016). Consequently, build-ups that involve non-spicular (“keratose”) demosponges might be much more widespread in time and space as currently realized (Szulc 1997; Bachmann 2002; Luo & Reitner 2014, 2016; Lee & Riding 2020). At the same time, the ecological meaning of microbe-metazoan build-ups remains somewhat elusive, particularly with regard to local and global environmental conditions.

The taxonomic term “Keratosa” was first introduced by Grant (1861) and further established by following work (e.g., Bowerbank, 1864; von Lendenfeld, 1889; Minchin, 1900). Later biochemical and molecular phylogenetic studies showed that these sponges belong to a sister group of the spicular demosponges that consists of two clades (e.g. Borchiellini et al. 2004; Lavrov et al. 2008; Erpenbeck et al. 2012). One clade, confusingly also termed “Keratosa”, is characterized by organic skeletons made of spongin, whilst the second order Verongimorpha exhibit chitinous skeletons (Morrow & Cárdenas 2015; Vacelet et al. 2019). Due to the strong homoplasy in the skeletal morphology of these sponges (e.g., Vacelet et al. 2019; Erpenbeck et al. 2020) and the loss of biochemical information during taphonomic processes, it is so far challenging to differentiate the two clades in the fossil record. The nomenclature is additionally complicated by the term “keratolite”. This term was introduced by Lee & Riding (2021) for associations of calcified non-spicular demosponges and microbialites; a relationship that was firstly described in detail by Luo & Reitner (2016). Problematically, however, the term “keratolite” literally means “horny skin rock” and is also used to refer to pathological skin structures, keratine faser proteins of various origin, and for various commercial products such as cosmetic and medical skin care. Thus, the term “keratolite” is misleading, although it might sound similar to “keratose”. In order to avoid confusion, we will use the term “non-spicular demosponges” for fossil materials.

Based on three-dimensional reconstruction of fossil specimens, fibrous/filamentous networks made of microspars correlate well with skeletal elements of modern non-spicular demosponges, made of spongin/chitin (Fig. 1; Luo & Reitner 2014; 2016). The clotted to peloidal micrite in between the networks correlates well with sponge tissue (Fig. 1). Later work applied “vermiform” fabric, namely ‘a pervasive meshwork of narrow anastomosing light-coloured, microspar-filled, tubules of varied diameter, ~20–50 μm wide’ to describe sponges (Lee & Riding 2020). Problematically, however, “vermiform” was formerly introduced to describe a distinctive microstructure in Phanerozoic stromatolites, such as *Madiganites mawsoni* and *Ilicta composite* Sidorov (Walter 1972). It consists of ‘narrow, sinuous areas of sparry carbonate surrounded by darker, usually finer-grained, carbonate’ (Walter 1972). To avoid potential confusion with the original definition of “vermiform” *sensu* Walter (1972), we here use “meshlike” fabric, which represents former organic skeleton of non-spicular demosponges (Fig. 1). The Neoproterozoic – early Cambrian interval shows a marked decline in stromatolite abundance. This development might be due to competition with eukaryotes, a reduced fossilization through authigenic minerals resulting from changing seawater chemistries, or a combination thereof (Grotzinger & Knoll 1999; Riding 2000; Arp et al. 2001). Whatever the reason is, in the Phanerozoic, stromatolites only developed in specific environments and/or during exceptional time intervals (Chen et al. 2019). For instance, stromatolites re-diversified profoundly in the aftermath of the Permian – Triassic Crisis – along with other types of microbialites and unusual sedimentary features (Woods 2014; Friesenbichler et al. 2018; Heindel et al. 2018; Chen et al. 2019; Pei et al. 2019). The widespread proliferation of microbial mats likely resulted from a suppressed ecological competition during this time (Foster et al. 2020). This situation supposedly lasted until “classic” metazoan ecosystems completely recovered and finally re-established during the mid–late Anisian, Middle Triassic, that is, 8-9 million years (Myr) after the crisis (Chen & Benton 2012). Thus, Lower to Middle Triassic “stromatolites” represent an important piece of the puzzle in understanding ecosystem recovery after the Permian – Triassic Crisis (Lehrmann 1999; Kershaw et al. 1999, 2012; Wang et al. 2005, 2019; Yang et al. 2011; Fang et al. 2017; Wu et al. 2017; Chen et al. 2019; Pei et al. 2019).

**Fig. 1.**
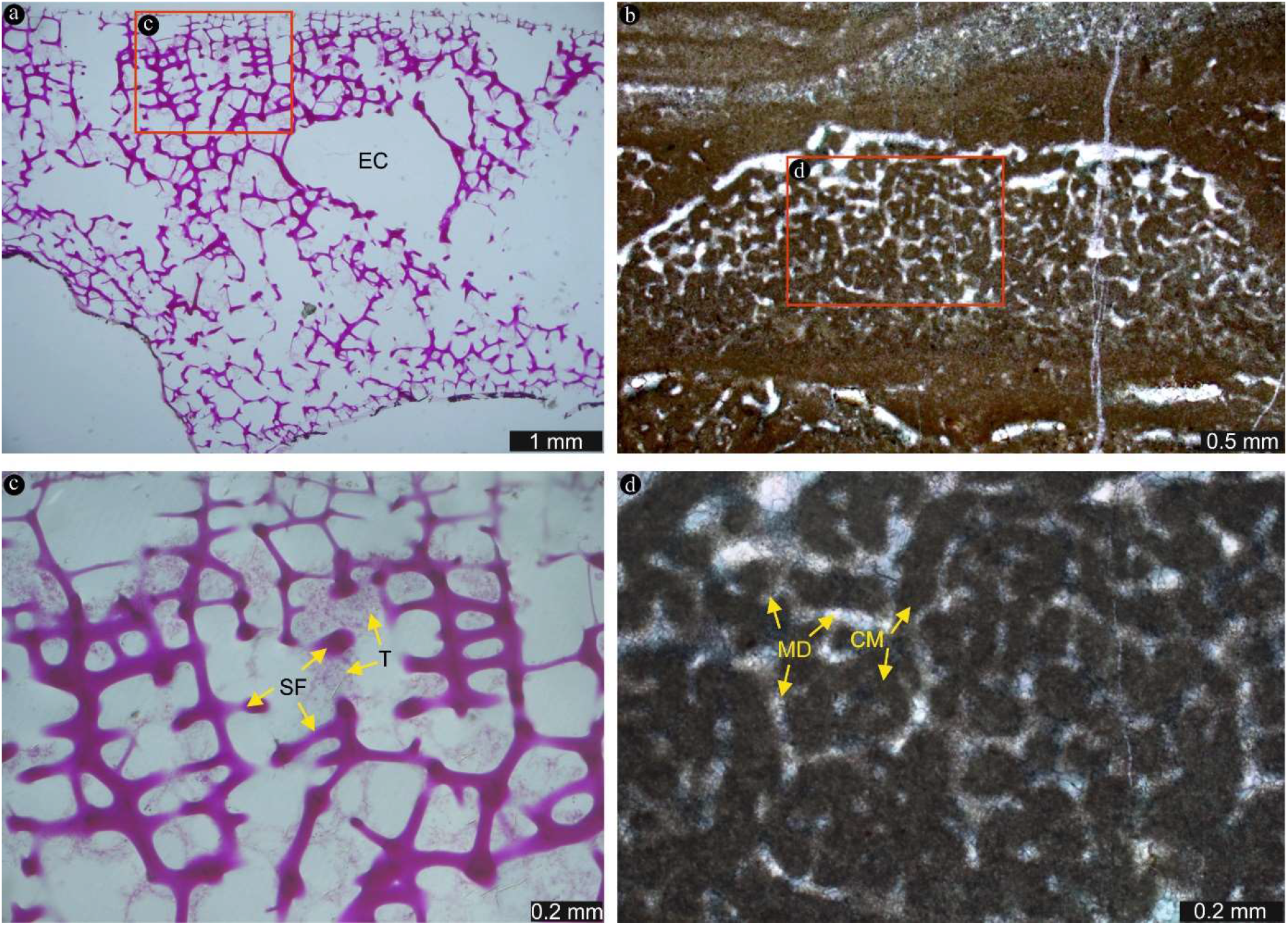
Comparison of dictyoceratid, modern non-spicular demosponge from a cave of Hicks Reef, Great Barrieer Reef, Australia with non-spicular demosponges in Auerstedt microbe-metazoan build-ups. The rectangle part in (a), stained with basic fuchsine, is the same as (c). The rectangle part in (b), stained with Alizarin Red S and K-hexacyanoferrate II in a 0.2 % HCl solution (cold), is the same as (d). Fibrous/filamentous networks made of spongin fibers (SF) (c) correlate well with microsparitic dolomite (MD) (d). The clotted to peloidal micrite (CM) correlates well with sponge tissue (T) (c). EC=Excurrent Canal

There are at least five marked stromatolite occurrences during Early to Middle Triassic times in the Germanic Basin: (1) the Lower Buntsandstein (Kalkowsky, 1908; Paul and Peryt, 2000; Paul et al. 2011), (2) the Upper Buntsandstein (Naumann 1928), (3) the lower Middle Muschelkalk (Hagdorn & Simon 1993), (4) the upper Middle Muschelkalk (Szulc 1997; Krause & Weller 2000; Luo & Reitner 2014), and (5) the Lower Keuper (Bachmann 2002; Luo & Reitner 2016). Despite their significance to understand ecosystem recovery after the Permian – Triassic Crisis, these microbialites have barely been studied from a geobiological perspective so far. For instance, there are very few studies on late Anisian (i.e., upper Middle Muschelkalk) “stromatolites” and associated facies in the Germanic Basin (Szulc 1997; Krause & Weller 2000; Luo & Reitner 2014). We here aim to fill this gap by investigating the geobiology of late Anisian “stromatolites” in the Germanic Basin in greater detail. More specifically, we show that these structures actually consist of interlayered microbial mats and metazoans such as non-spicular demosponges, and therefore should more correctly be referred to as microbe-metazoan build-ups. Furthermore, we discuss paleoecological implications of these build-ups, particularly with regard to ecosystem recovery after the Permian – Triassic Crisis.

### Geological setting

The Germanic Basin covered large parts of central Europe during the Permian and Triassic periods (Fig. 2). Paleogeographically, it was a peripheral basin of the western Tethys Ocean (Ziegler 1990) characterized by a subtropical climate (Scotese & McKerrow 1990). During the Anisian (Middle Triassic), the Germanic Basin and the Tethys were connected by three gates, namely (from northeast to southwest) the East Carpathian Gate, the Silesian – Moravian Gate, and the Western Gate (Götz & Feist-Burkhardt 2012; Ziegler 1990). The sites investigated in this study were located in the northeastern and southern parts of the Germanic Basin (Auerstedt, Mauer and Seyweiler, respectively), and close to the Silesian – Moravian Gate (Libiąż) (Fig. 2).

**Fig. 2.**
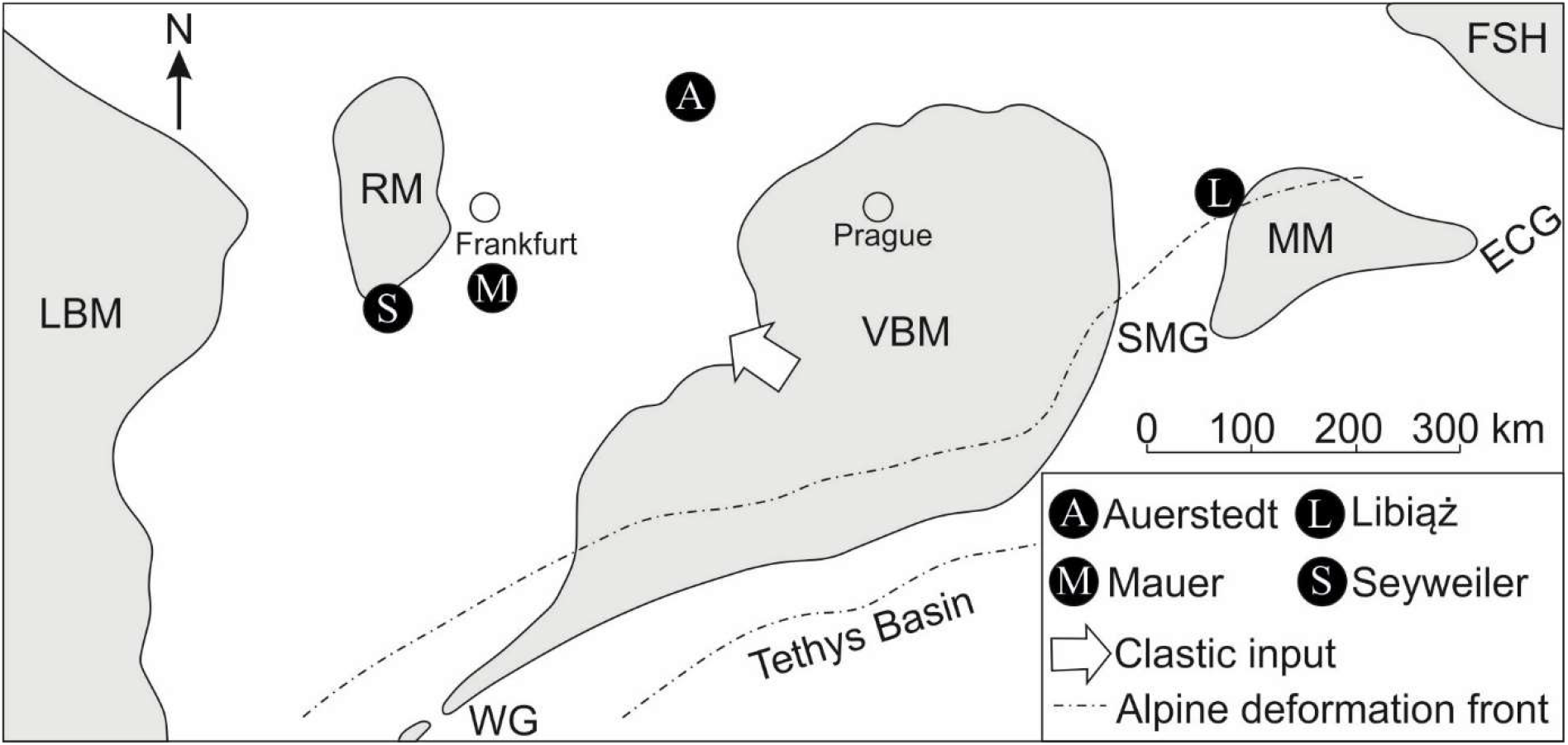
Paleogeographic map of the Germanic Basin during Permian and Triassic times. During the Anisian (i.e., Middle Triassic), the Germanic Basin and the Tethys were connected by three gates, namely (from northeast to southwest) the East Carpathian Gate (ECG), the Silesian – Moravian Gate (SMG), and the Western Gate (WG) (Götz & Feist-Burkhardt 2012; Ziegler 1990). The investigated outcrops are located in the northern (A: Auerstedt) and southern part of the Germanic Basin (M: Mauer and S: Seyweiler) as well as close to the Silesian – Moravian Gate (L: Libiąż). LBM: London – Brabant – Massif, RM: Rhenish Massif, VBM: Vindelician – Bohemian – Massif, MM: Malopolska Massif, FSH: Fenno – Scandian High

Three of the outcrops (Auerstedt, Mauer and Seyweiler) can be correlated by a chert layer (Urlichs 1992). Sedimentary rocks exposed at these localities belong to the Diemel Formation, which consists of grey–yellow dolomite, dolomitic limestone and marls (Hagdorn & Simon 2005; Hagdorn 2010) (Fig. 3). The formation locally contains ooids and is characterized by an euryhaline, species-poor fauna and stromatolites (Hagdorn & Simon, 2005). At Libiąż, stromatolite-bearing intervals occur in the upper part of the *Diplopora* Beds (Szulc 1997). Since both the Diemel Formation and the *Diplopora* Beds are of Illyrian age (i.e., late Anisian, Middle Triassic) (Szulc 1997; Hagdorn & Simon 2005; Hagdorn 2010), the investigated “stromatolites” are approximately coeval.

**Fig. 3.**
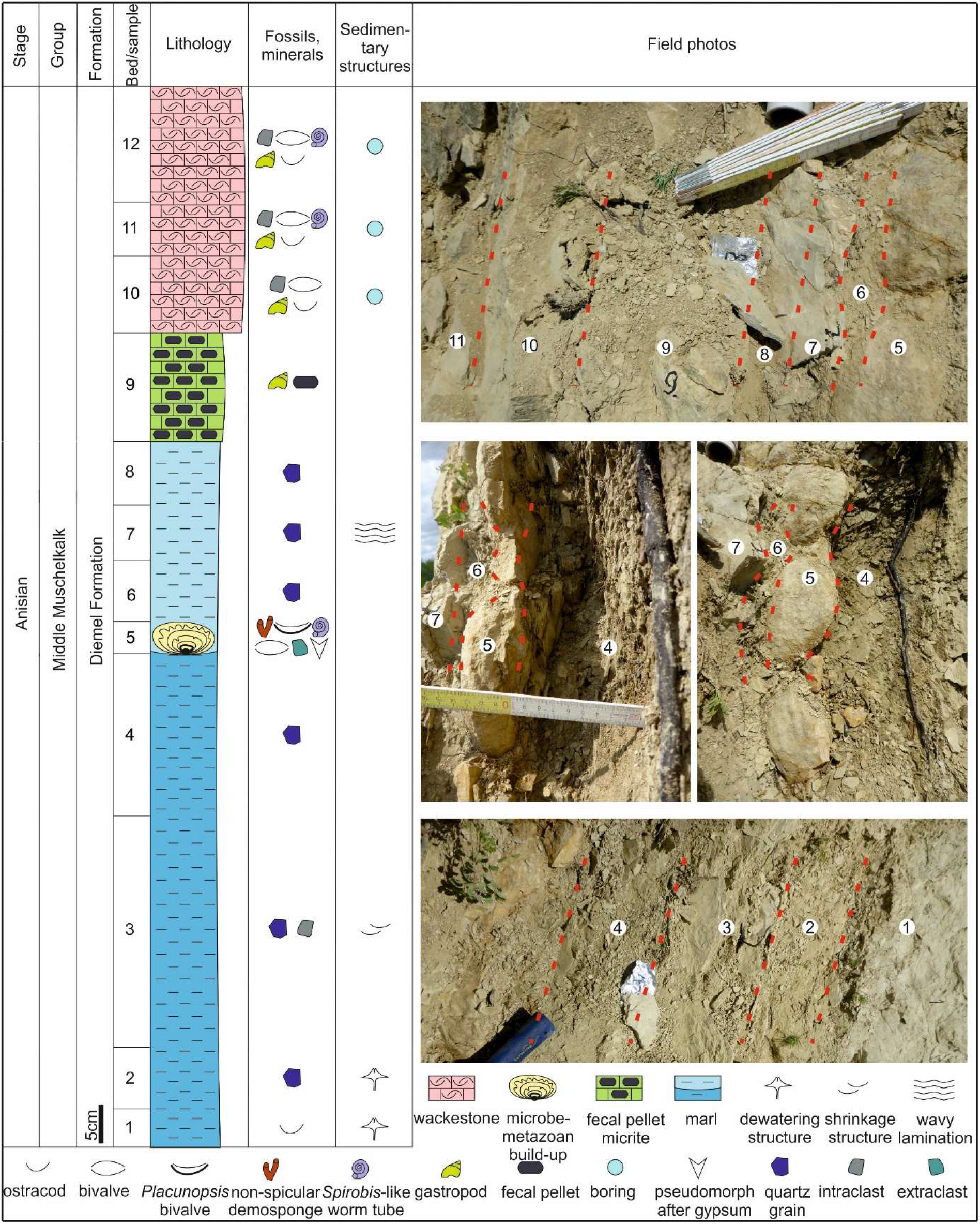
The Auerstedt section, showing stratigraphical, sedimentological and paleontological features in late Anisian. Numbers in the field photos correspond to those in bed/sample column

## Materials and methods

### Fieldwork and petrography

Fresh samples from Auerstedt (microbe-metazoan build-ups and associated facies) as well as from Mauer, Seyweiler and Libiąż (microbe-metazoan build-ups) were taken.

Petrographic thin sections were prepared and analyzed using a Zeiss SteREO Discovery.V12 stereomicroscope and a Zeiss AXIO Imager.Z1 microscope. In each case, photos were taken with an AxioCamMRc 5 MB camera.

### Analytical imaging technique

Micro X-ray fluorescence (μ-XRF) was applied to obtain element distribution images of thin sections. The analyses were conducted with a Bruker M4 Tornado instrument equipped with an XFlash 430 Silicon Drift Detector. Measurements (spatial resolution = 70 μm, pixel time = 30 ms) were performed at 50 kV and 400 μA at a chamber pressure of 20 mbar.

Raman single spectra and two-dimensional spectral images were collected using a WITec alpha300R fiber-coupled ultra-high throughput spectrometer. Before analysis, the system was calibrated using an integrated light source. For Raman single spectra, the experimental setup includes a 405 nm excitation laser (chosen to reduce fluorescence effects), 10 mW laser power, a 50× long working distance objective with a numerical aperture of 0.55, and a 1200 g mm^−1^ grating. The spectrometer was centered at 1530 cm^−1^, covering a spectral range from 122 cm^−1^ to 2759 cm^−1^. This setup has a spectral resolution of 2.6 cm^−1^. Each spectrum was collected by two accumulations, with an acquisition time of 5 s. For Raman spectral images, the experimental setup includes a 532 nm excitation laser, an automatically controlled laser power of 30 mW, a 50× long working distance objective with a numerical aperture of 0.55, and a 300 g mm^−1^grating. The spectrometer was centered at 2220 cm^−1^, covering a spectral range from 68 cm^−1^ to 3914 cm^−1^. This setup has a spectral resolution of 2.2 cm^−1^. Spectra were collected at a step size of 1 μm in horizontal and vertical direction by an acquisition time of 100 ms for each spectrum. Automated cosmic ray correction, background subtraction, and fitting using a Lorentz function were performed using the WITec Project software. Raman images were additionally processed by spectral averaging/smoothing and component analysis.

### Bulk analysis

Total organic carbon (TOC) analysis and Rock-Eval pyrolysis of two samples of Auerstedt and Mauer build-ups were performed at Applied Petroleum Technology. TOC measurements were executed using a Leco SC-632 instrument. Rock-Eval pyrolysis was conducted using a HAWK instrument and followed established protocols. Briefly, the temperature was ramped from 300 °C (held for 3 min) to 650 °C at 25 °C min^−1^. The measurements included quantities of free hydrocarbons (S1, mg HC /g rock), hydrocarbons yielded from labile kerogen (S2, mg HC /g rock), temperatures at maximum yields of S2 hydrocarbons (*T_max_*, °C), as well as the CO_2_ generated from organic carbon at higher temperature up to 650 °C (S3, mg CO_2_ /g rock). These data were used to calculate the hydrogen index (HI; S2 /TOC*100), the oxygen index (OI; S3 /TOC*100) and the production index (PI; S1 /(S1+S2)).

### Sedimentary hydrocarbons preparation and analysis

6 samples from Auerstedt (1 build-up and 4 associated facies) and Mauer (1 build-up) were prepared for sedimentary hydrocarbon analyses using established methodology (Duda 2014; Duda et al. 2014a, b, 2020). All laboratory materials used were heated to 500 °C for 3 h and/or extensively rinsed with acetone. Only distilled solvent was used for extraction and further work. Additionally, a blank sample (pre-combusted sea sand) was prepared and analyzed in parallel to keep track of laboratory contamination.

Sample surfaces were removed using a rock saw. The remaining blocks were crushed and powdered with a Retsch MM 301 pebble mill. Sample powders (25 g) were extracted with ~25 ml dichloromethane/methanol (DCM/MeOH; 9/1; v/v), ~25 ml DCM/*n*-hexane (DCM/*n*-hexane; 1/1; v/v), and ~25 ml n-hexane using ultrasonication (20 °C, 10 min). The resulting total organic extracts (TOEs) were concentrated using a rotary evaporator and a gentle stream of N2. Activated copper was added to remove elemental sulfur before the TOEs were analyzed by Gas chromatography – mass spectrometry (GC-MS).

GC-MS analysis was carried out with a Thermo Scientific Trace 1300 Series GC coupled to a Thermo Scientific Quantum XLS Ultra MS. The GC was equipped with a capillary column (Phenomenex Zebron ZB-5, 30 m, 0.25 μm film thickness, 0.25 mm inner diameter). TOEs were injected into a splitless injector and transferred to the GC column at 300 °C. Helium was used as carrier gas with a constant flow rate of 1.5 ml min^−1^. The GC oven temperature was held isothermal at 80 °C for 1 min and then ramped to 310 °C at 5 °C min^−1^, at which it was kept for 20 min. Electron ionization mass spectra were recorded in full scan mode (electron energy = 70 eV; mass range = *m/z* 50 – 600, scan time = 0.42 s).

### Stable isotope analysis (δ^13^C_carb_, δ^18^O_carb_)

43 samples (ca. 100μg each) were taken with a high-precision drill from individual mineral phases of polished rock slabs. Additionally, 11 fresh (i.e. not weathered) bulk rock samples were crushed with a hammer and/or powdered with a Retsch MM 301 pebble mill. In these cases, approximately 100 - 200 μg were used for bulk analyses. The measurements were performed at 70 °C using a Thermo Scientific Kiel IV carbonate device coupled to a Finnigan DeltaPlus gas isotope mass spectrometer. Carbon and oxygen stable isotope ratios of carbonate minerals are reported as delta values (δ^13^C_carb_ and δ^18^O_carb_) relative to Vienna Pee Dee Belemnite (VPDB) reference standard. Reproducibility was confirmed to be <0.1 by replicate analyses of a standard (NBS19).

All preparation and analytical work except for bulk analyses (see 3.3) have been carried out at the Geoscience Center of the Georg-August-Universität Göttingen.

## Results

### Microbe-metazoan build-ups and associated facies

#### Sedimentary succession at Auerstedt

The lower part of the Auerstedt section is dominated by marls (beds 1–4, 6–8: Fig. 3). Microbe-metazoan build-ups are restricted to one distinct bed within this interval (i.e., bed 5). The marls below the microbe-metazoan build-ups show abundant dewatering and shrinkage structures (beds 1–2 and bed 3, respectively: Figs. 3; 4a), but these structures disappear section upwards. Above the microbe-metazoan build-ups, marls locally show wavy lamination structures (bed 7: Fig. 4b).

**Fig. 4.**
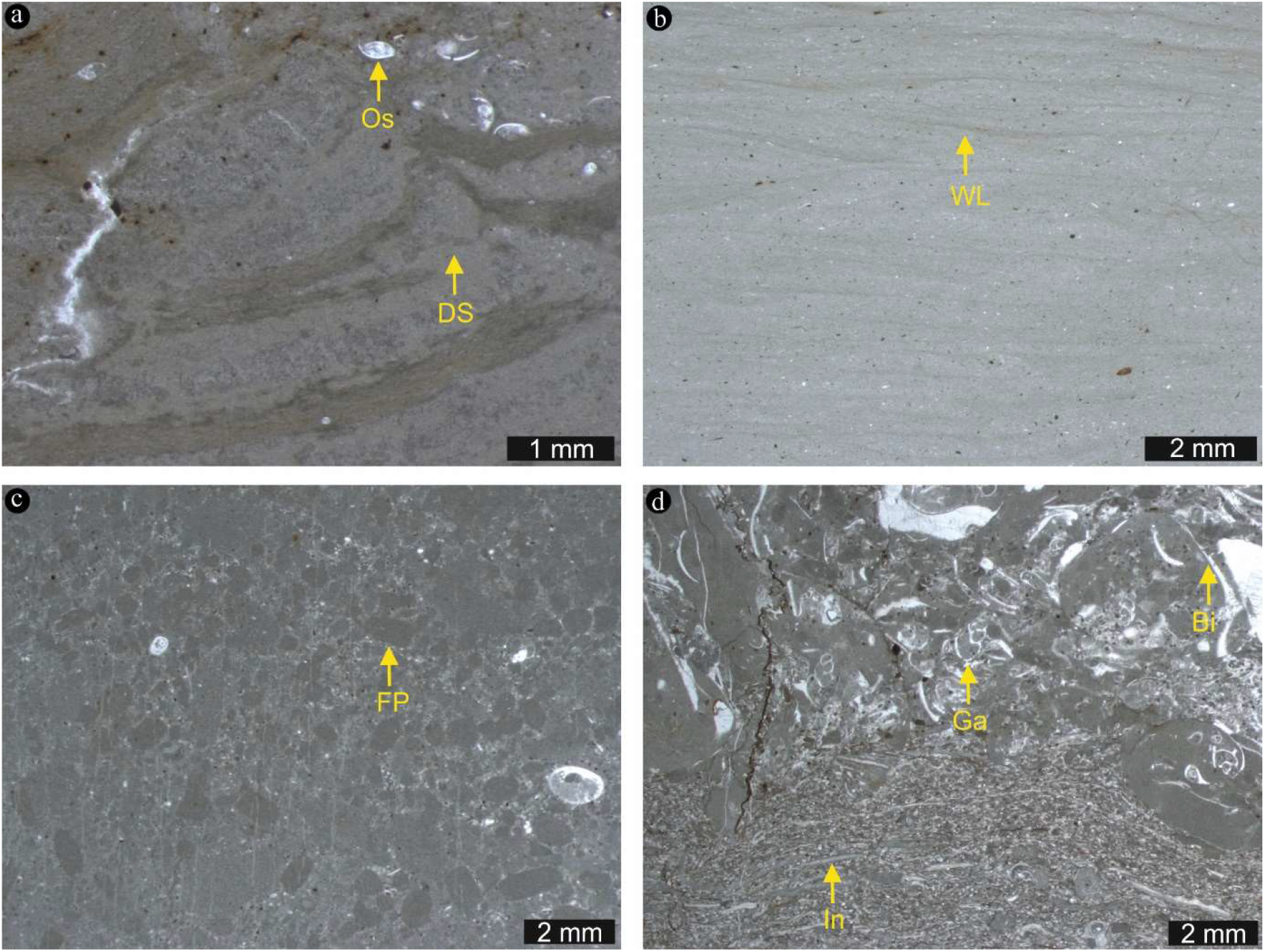
Thin section images (transmitted light) of strata below and above the Auerstedt microbe-metazoan build-ups. (a) The marls below the microbe-metazoan build-ups show abundant dewatering structures (DS). Marls below and above the microbe-metazoan build-ups are generally fossils poor and contain only a few ostracods (Os). (b) Marls above the microbe-metazoan build-ups locally show wavy lamination (WL). (c) The marl interval is overlain by a micrite bed that contains abundant fecal pellets of crustaceans (FP). (d) Wackestone from the top of the section contains abundant intraclasts (In) and fossils, including ostracods, bivalves (Bi) and gastropods (Ga)

Overall, the marls contain abundant angular quartz grains and a few intraclasts (Fig. 3). They are generally poor in fossils and locally only contain few ostracods (Figs. 3; 4a). All other body fossils in this interval are directly associated with the microbe-metazoan build-ups that occur in bed 5 (see 4.1.2). The marl interval is overlain by a micrite bed that contains abundant fecal pellets of crustaceans (bed 9: Figs. 3, 4c). This unit is followed by wackestone which contains abundant intraclasts and fossils (beds 10–12: Figs. 3; 4d). Fossils in this unit include ostracods, gastropods and bivalves, the latter commonly showing borings (Figs. 3; 4d).

#### Auerstedt microbe-metazoan build-ups

Microbe-metazoan build-ups at Auerstedt are limited to bed 5 (Fig. 3). The build-ups are about 5 cm thick and exhibit flat-domal shapes. Individual domal bodies intergrow laterally, resulting in a horizontally continuous bed. About 0.5 cm thick, round-shaped mud chips in the top part of the underlying marl layer (bed 4; Fig. 3) might have served as stable substrates for biological communities that formed the build-ups (Fig. 5). Some mud chips are well preserved (Fig. 5a), whilst others are not (Fig. 5b). The biocrusts at the top of the build-ups consist of encrusting *Placunopsis* bivalves and *Spirobis-like* worm tubes (Fig. 6).

**Fig. 5.**
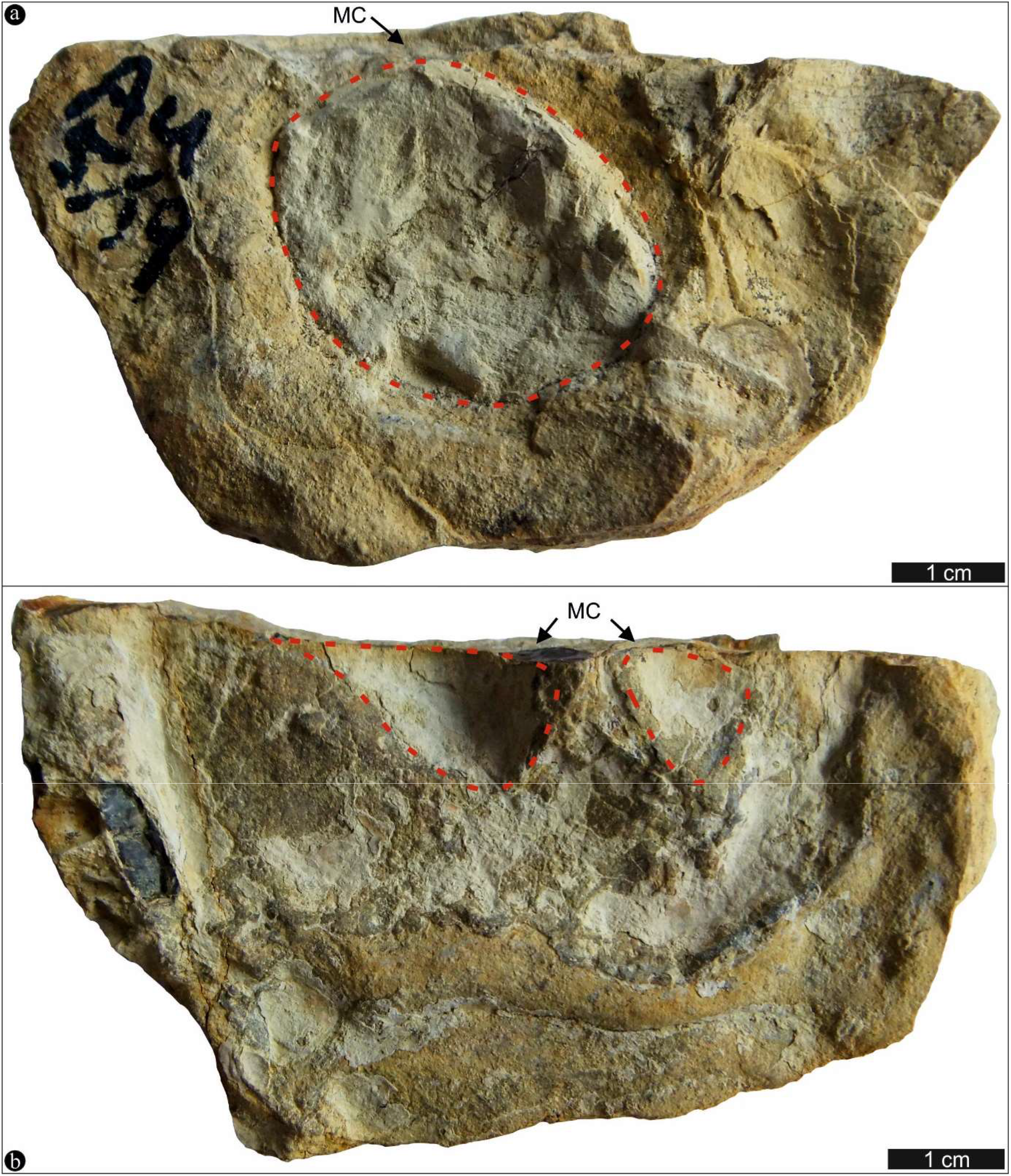
Photos of the lower surface of microbe-metazoan build-ups from Auerstedt. Note the round-shaped mud chips (MC) that likely served as stable substrates for the growth of the build-ups. Some mud chips are well preserved (a), whilst others are not (b)

**Fig. 6.**
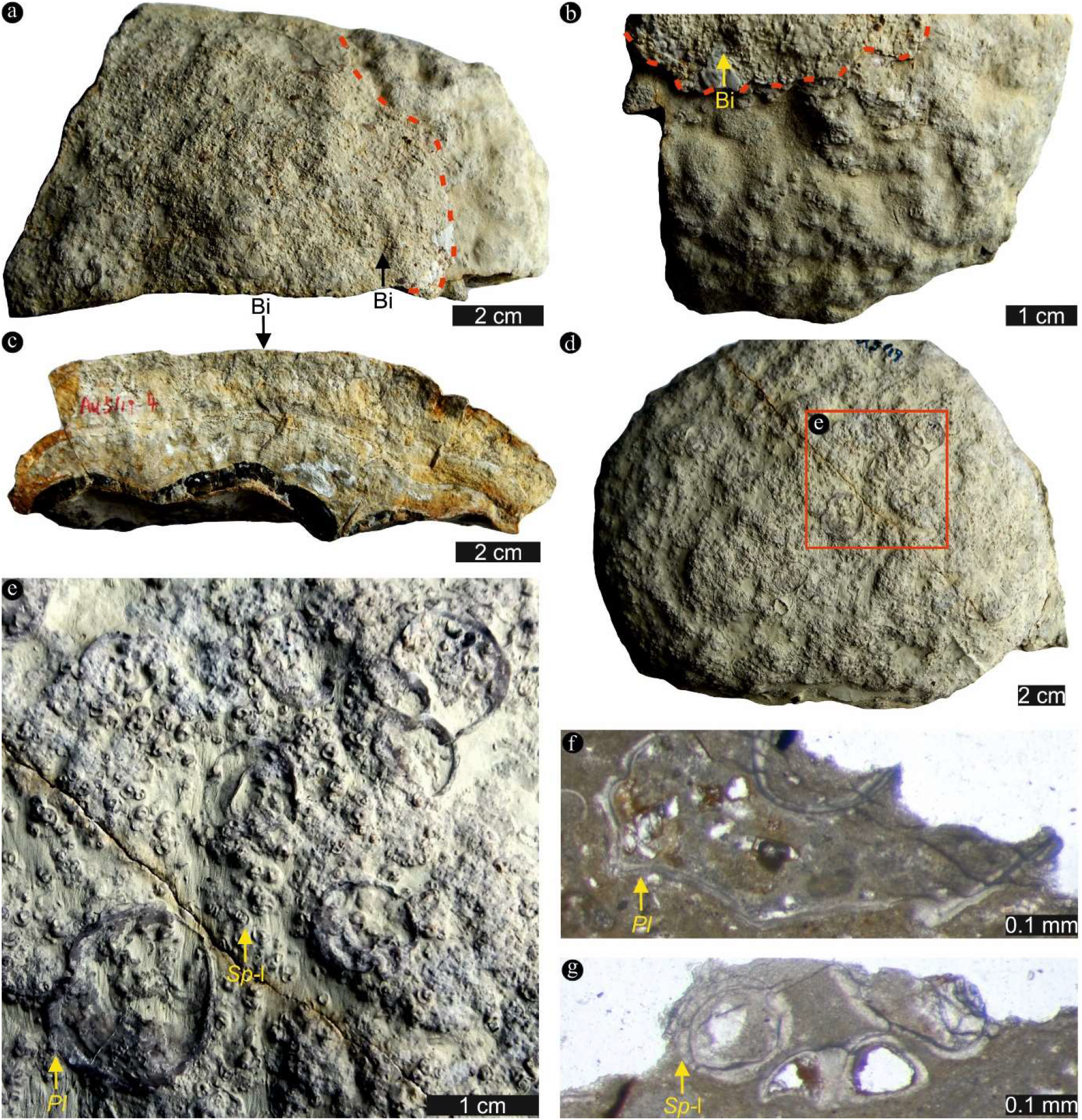
Photos and thin section images of biocrusts (Bi) at the top of microbe-metazoan buildups from Auerstedt (a–e and f, g, respectively). (e) is a close-up of the rectangle part in (d). These biocrusts are formed by *Placunopsis* bivalves *(Pl)* and *Spirobis-like* worm tubes (*Sp-*l)

The microbe-metazoan build-ups exhibit a complex architecture with up to 13 layers (Fig. 7a). The layers are mainly comprised of calcite, dolomite, silica and organic matter as revealed by μ-XRF and Raman spectroscopy (single spectra and spectral images) (Figs. 8, 9). Layers 1–3 are in total about 0.7 cm thick (Fig. 7a). Layers 1 and 3 consist of non-laminated micritic calcite and dolomite (Figs. 8b, e; 10a). Layer 2 is characterized by a similar mineralogy, but additionally exhibits microbial lamination features and contains some bivalves (Fig. 10a). Layer 4 is about 0.7 cm thick (Fig. 7a). It is composed of quartz (chert) as confirmed by relative Ca depletions as well as Si enrichments in μ-XRF images and Raman spectroscopy (Fig. 8b, c, f). However, the silicification is not fabric destructive since planar laminations can still be observed (Fig. 10b).

**Fig. 7.**
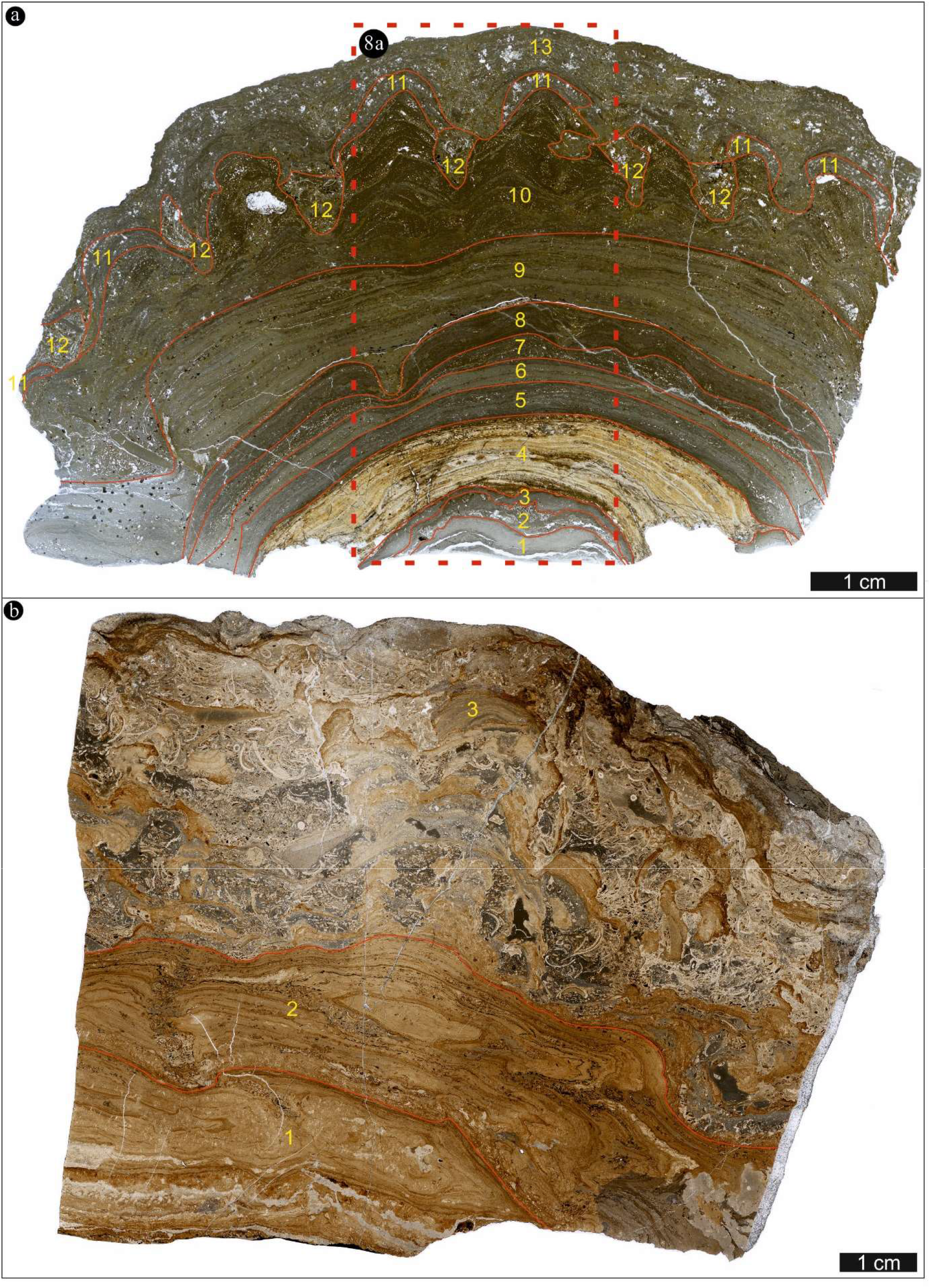
Scan images (transmitted light) of microbe-metazoan build-ups from Auerstedt (a) and Mauer (b). The rectangle dashed line in (a) indicates the location of Fig. 8a

**Fig. 8.**
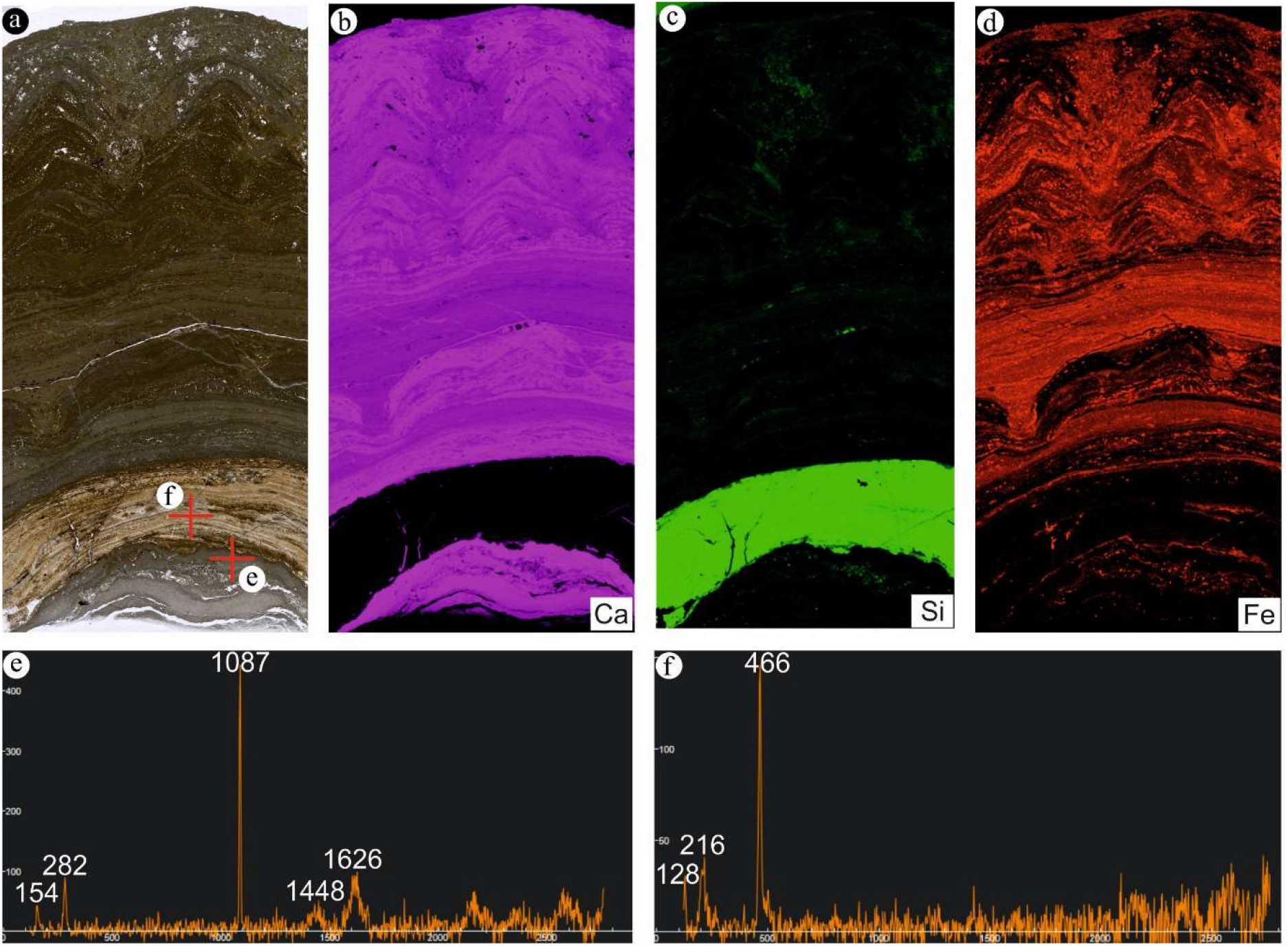
Micro X-ray fluorescence (μ-XRF) images and Raman spectroscopic features of Auerstedt microbe-metazoan build-ups. (a) Scan image (transmitted light). (b) Calcium distribution. (c) Silicon distribution. (d) Iron distribution. (e) Raman spectrum of calcite (location shown in (a)). (f) Raman spectrum of quartz (chert) (location shown in (a))

**Fig. 9.**
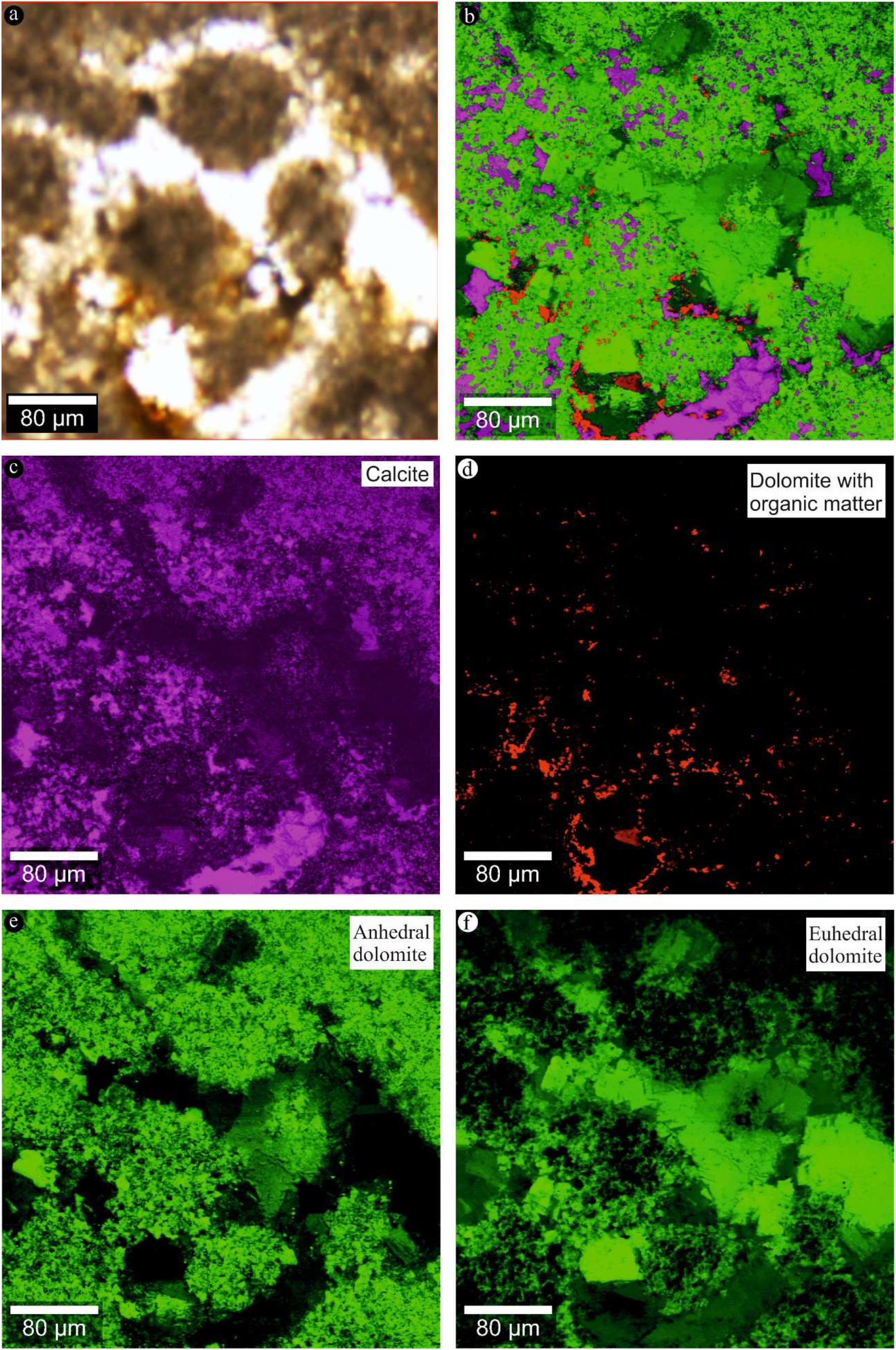
Raman spectral images of Auerstedt microbe-metazoan build-ups. (a) Thin section image (transmitted light). (b) Combined image of calcite, dolomite and organic matter. (c) Distribution of calcite. (d) Distribution of dolomite with organic matter. (e–f) Distribution of anhedral and euhedral dolomite, respectively. Colour code: purple = calcite, red = dolomite with organic matter, green = dolomite

**Fig. 10.**
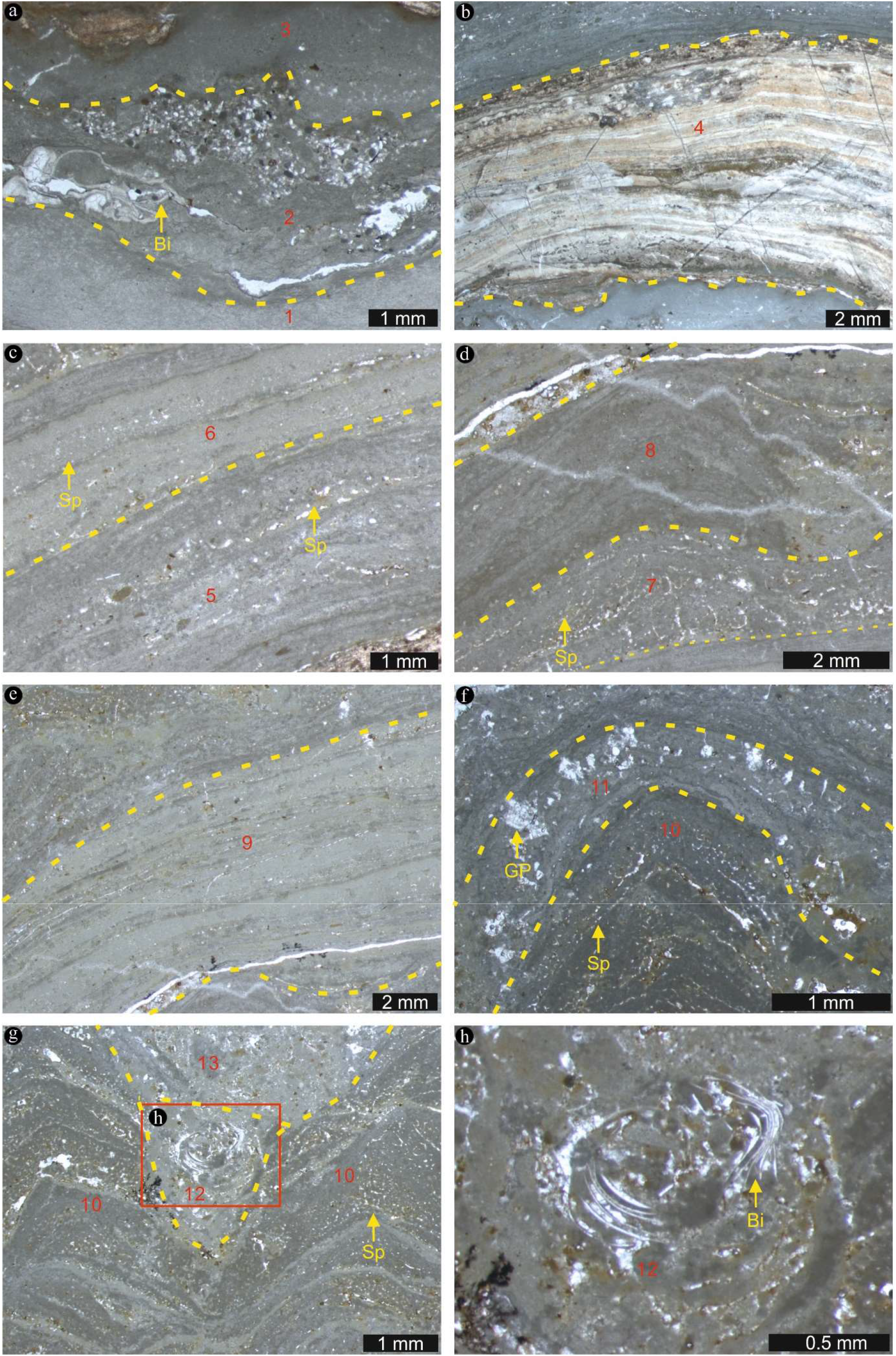
Thin section images of microbe-metazoan build-ups from Auerstedt. (a) Layers 1 and 3 both consist of non-laminated micritic calcite and dolomite. Layer 2 is characterized by a similar mineralogy, but additionally exhibits microbial lamination features and contains some bivalves (Bi). (b) Layer 4 is composed of quartz (chert) and shows planar laminations. (c) Layers 5 and 6 consist of planarly laminated micritic calcite and dolomite. Both layers locally exhibit clotted to peloidal features, as well as mesh-like fabrics which is more dominant in layer 5 as compared to layer 6. (d) Layers 7 and 8 are composed of clotted to peloidal micritic calcite and dolomite that exhibit wavy to domal laminations. Notably, layer 7 contains sponges, showing dense mesh-like fabrics. (e) Layer 9 consists of planarly laminated micritic calcite and dolomite, locally exhibiting a peloidal feature. (f) Layer 10 consists of dense clotted to peloidal micritic calcite and dolomite. It exhibits domal to conical laminations and mesh-like fabrics. The overlying layer 11 comprises non-laminated micritic calcite and dolomite and contains abundant calcite pseudomorphs after gypsum (GP). (g–h) Layer 12 encompasses the interspaces between laminated cones, magnified in (h). These interspaces filled with abundant fossils of bivalves and ostracods, resulting in a wackestone fabric. Layer 13 consists of non-laminated micritic calcite and dolomite and contains abundant calcite pseudomorphs after gypsum. Sp = sponge

The younger layers 5 and 6 have a total thickness of ~0.4 cm (Fig. 7a) and consist of planarly laminated micritic calcite and dolomite (Fig. 10c). Both layers locally exhibit clotted to peloidal features, as well as mesh-like fabrics which is more dominant in layer 5 as compared to layer 6 (Fig. 10c). Layers 7 and 8 have a total thickness of ~0.4 cm (Fig. 7a) and are composed of clotted to peloidal micritic calcite and dolomite that exhibit wavy to domal laminae (Fig. 10d). Notably, layer 7 contains sponge fossils that show dense mesh-like fabrics (Fig. 10d). Layer 9 has a thickness of ~0.6 cm (Fig. 7a) and consists of planarly laminated micritic calcite and dolomite (Fig. 10e). Its clotted to peloidal fabric is less distinct than that of layer 5 but more explicit than that of layer 6 (Fig. 10c). Layer 10 is ~1.6 cm thick (Fig. 7a) and consists of dense clotted to peloidal micritic calcite and dolomite (Fig. 10f). The layer exhibits domal to conical laminations and a mesh-like fabric (Fig. 10f). The overlying layer 11 is ~0.2 cm thick (Fig. 7a), comprises non-laminated micritic calcite and dolomite, and contains abundant calcite pseudomorphs after gypsum (Fig. 10f). Layer 12 encompasses the interspaces between laminated cones (Fig. 7a). These interspaces exhibit a non-laminated wackestone texture and contain abundant bivalves and ostracods (Fig. 10g, h). Layer 13 is ~0.4 cm thick (Fig. 7a) and similar to layer 11 in lithology (Fig. 10g).

#### Microbe-metazoan build-ups from Mauer, Seyweiler and Libiąż

The investigated build-up from Mauer is ~8 cm thick and divided into three layers (Fig. 7b). The layers mainly consist of silicified carbonate as indicated by co-enrichments of Ca and Si observed in μ-XRF images. Layer 1 exhibits wavy laminations and is devoid of fossils (Fig. 11a). The following layer 2 shows wavy laminae with local interspaces, the latter containing abundant bivalves and ooids (Fig. 11b). In Layer 3, primary laminated fabrics appear to be strongly disturbed (Fig. 11c). Resulting sedimentary interspaces are filled with abundant bivalves, sponges, gastropods and ooids (Fig. 11c, d). Notably, bivalves are poorly sorted and randomly oriented (Fig. 11c). Sponge fossils display mesh-like fabrics, similar to those observed at Auerstedt (Fig. 11d).

**Fig. 11.**
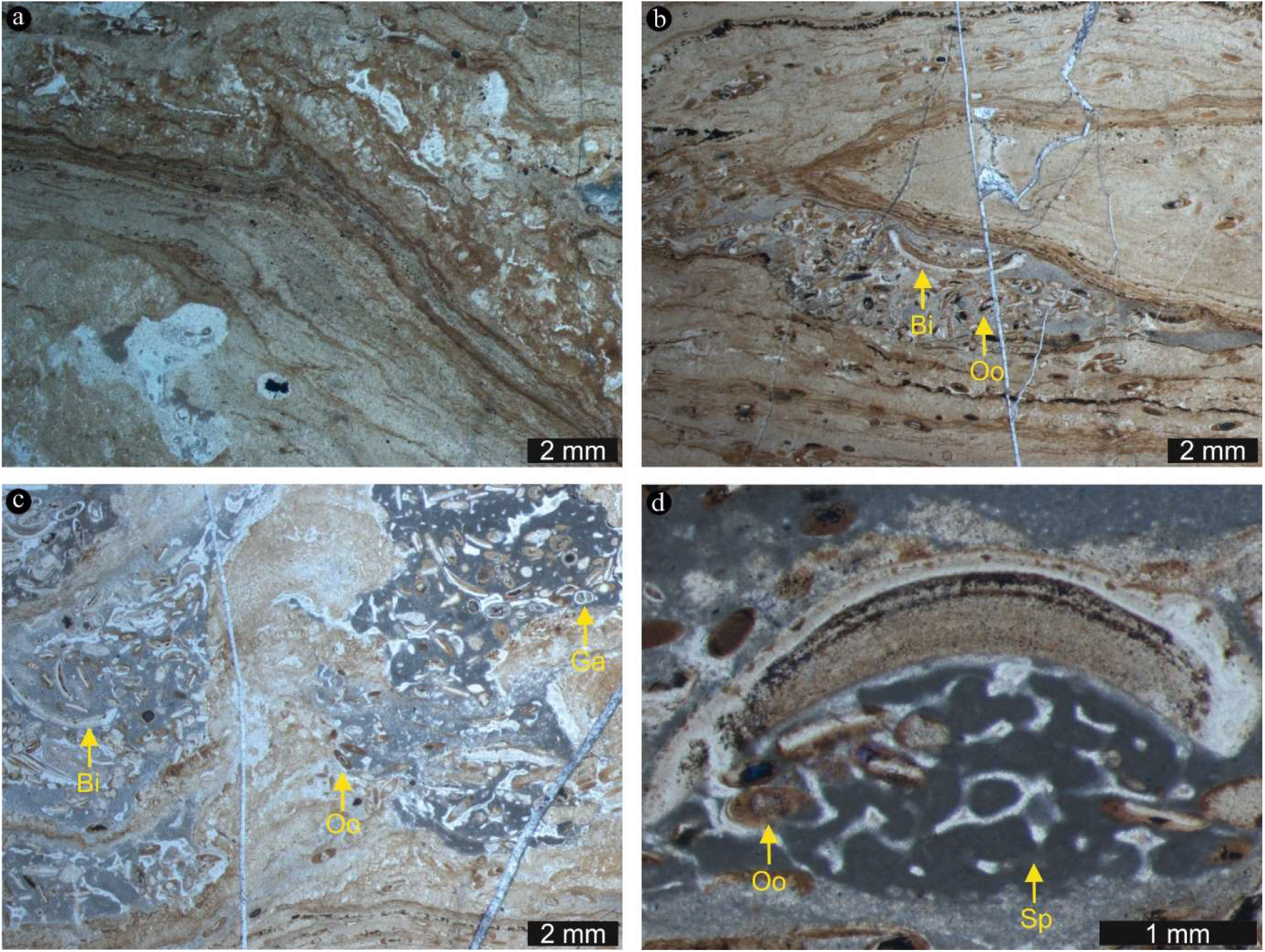
Thin section images of microbe-metazoan build-ups from Mauer. (a) Layer 1 is mainly made of silicified carbonate and exhibits wavy laminations. (b) Layer 2 also consists of silicified carbonate that shows wavy laminae. In contrast to layer 1, it contains abundant bivalves (Bi) and ooids (Oo). (c–d) Layer 3 also consists of silicified carbonate. Primary laminated fabrics appear to be strongly disturbed, and resulting sedimentary interspaces are filled with abundant bivalves, sponges (Sp), gastropods (Ga) and ooids. Notably, bivalves are poorly sorted and randomly oriented. Sponges display mesh-like fabrics

The studied microbe-metazoan build-up from Seyweiler is ~8 cm thick and mainly composed of silicified carbonate as shown by relative Ca and Si enrichments in μ-XRF images. The layers exhibit wavy and conical laminae (Fig. 12a–c) and locally enclose ooids (Fig. 12b). Fossils include bivalves and sponges (Fig. 12b–d). Some bivalves are vertically oriented (Fig. 12b). Sponges exhibit mesh-like fabrics as well (Fig. 12c, d).

**Fig. 12.**
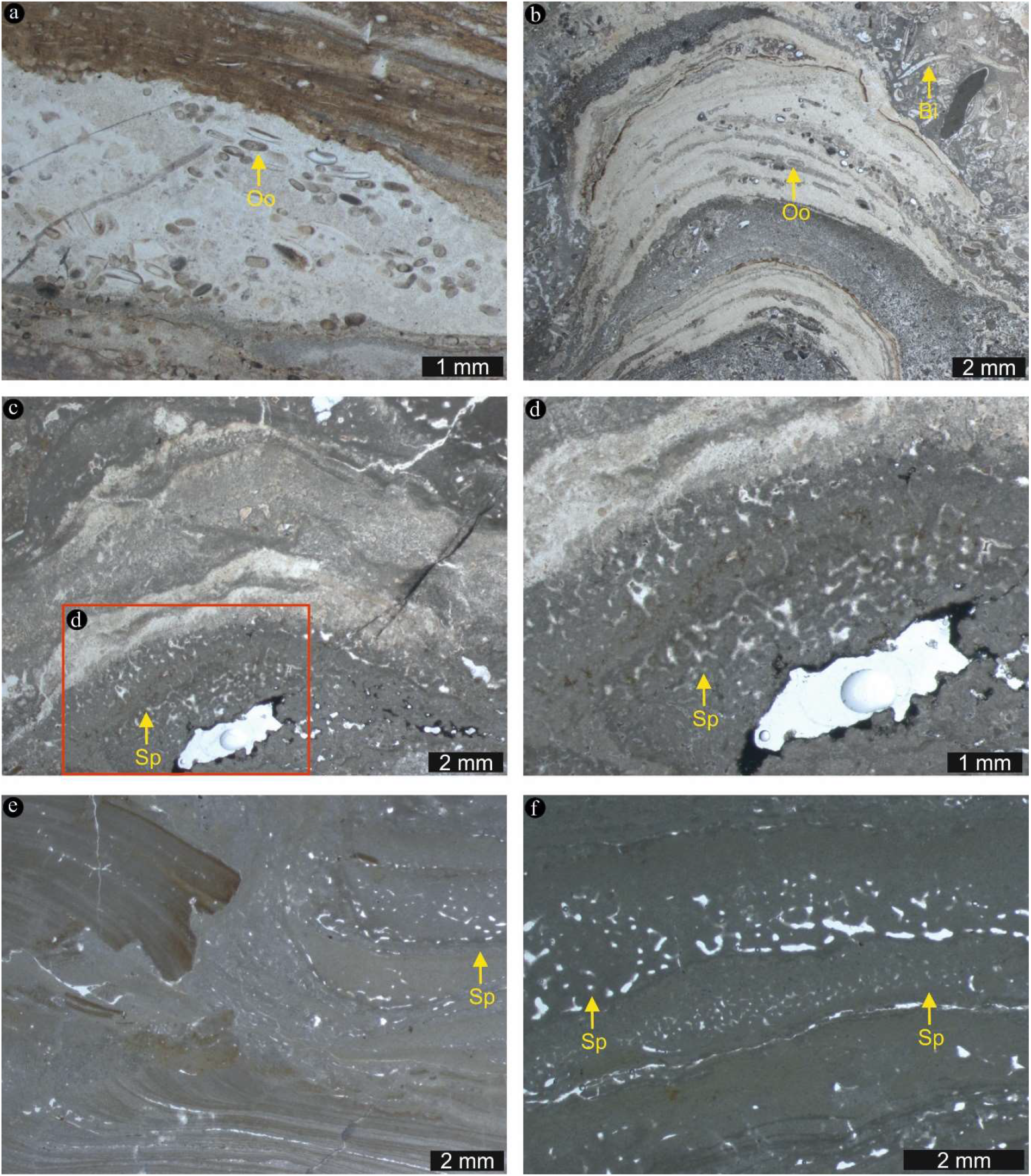
Thin section images of microbe-metazoan build-ups from Seyweiler and Libiąż. (a–d) Microbe-metazoan build-ups from Seyweiler consist of silicified carbonate and typically exhibit wavy and conical laminae (a–c). The conical laminae locally enclose ooids (b). Fossils include bivalves (Bi) and sponges (Sp). Some bivalves are vertically oriented (b). Sponges exhibit mesh-like fabrics (c, d). (e–f) At Libiąż, microbe-metazoan build-ups are mainly made of calcite, characterized by planar and sometimes domal laminations. The laminations are sometimes disrupted and/or fragmented. Sponges (Sp) are abundant and can be clearly recognized based on distinctive features such as mesh-like fabrics

The analyzed microbe-metazoan build-ups from Libiąż can be up to 40 cm thick (Luo & Reitner 2014; this study). They are mainly composed of calcite, characterized by planar and sometimes domal laminations (Fig. 12e). The laminations are sometimes disrupted and/or fragmented (Fig. 12e). Sponges are abundant and can be clearly recognized based on distinctive features such as mesh-like fabrics (Fig. 12f).

### Bulk organic matter and sedimentary hydrocarbons

The analyzed materials are generally relatively organic lean. For instance, samples from below and above the build-ups at Auerstedt were devoid of any GC-MS amenable sedimentary hydrocarbons. Only the build-up samples preserved detectable amounts of organic matter (Mauer: 0.14 wt.% TOC, Auer: 0.16 wt.% TOC: see Tab. 1). These samples also contained sedimentary hydrocarbons such as *n*-alkanes, terminally branched alkanes, and acyclic isoprenoids (norpristane, pristane, phytane) (Fig. 13). Unfortunately, these records could not be verified by means of Rock-Eval since TOC contents are required to ≥0.2 wt.% and S2 values are >0.5 mg HC g^−1^ rock (see Peters 1986; Peters & Cassa 1994) (Tab. 1). Therefore, Rock-Eval data has to be considered unreliable and must be discounted.

**Table 1.** Total organic carbon (TOC) contents and Rock-Eval pyrolysis parameters for 2 samples of Auerstedt and Mauer microbe-metazoan build-ups

| Sample | TOC (wt. %) | T<sub>max</sub> (°C) | S1 (mg HC /g rock) | S2 (mg HC /g rock) | S3 (mg CO<sub>2</sub>/g rock) | PI (S1/(S1+S2)) | HI (mg HC/g TOC) | OI (mg CO<sub>2</sub>/g TOC) |
| --- | --- | --- | --- | --- | --- | --- | --- | --- |
| Mauer microbe-metazoan build-up | 0.14 | 525 | 0.02 | 0.05 | 0.11 | 0.29 | 35 | 78 |
| Auerstedt microbe-metazoan build-up | 0.16 | 613 | 0.02 | 0.03 | 0.13 | 0.40 | 19 | 80 |

**Fig. 13.**
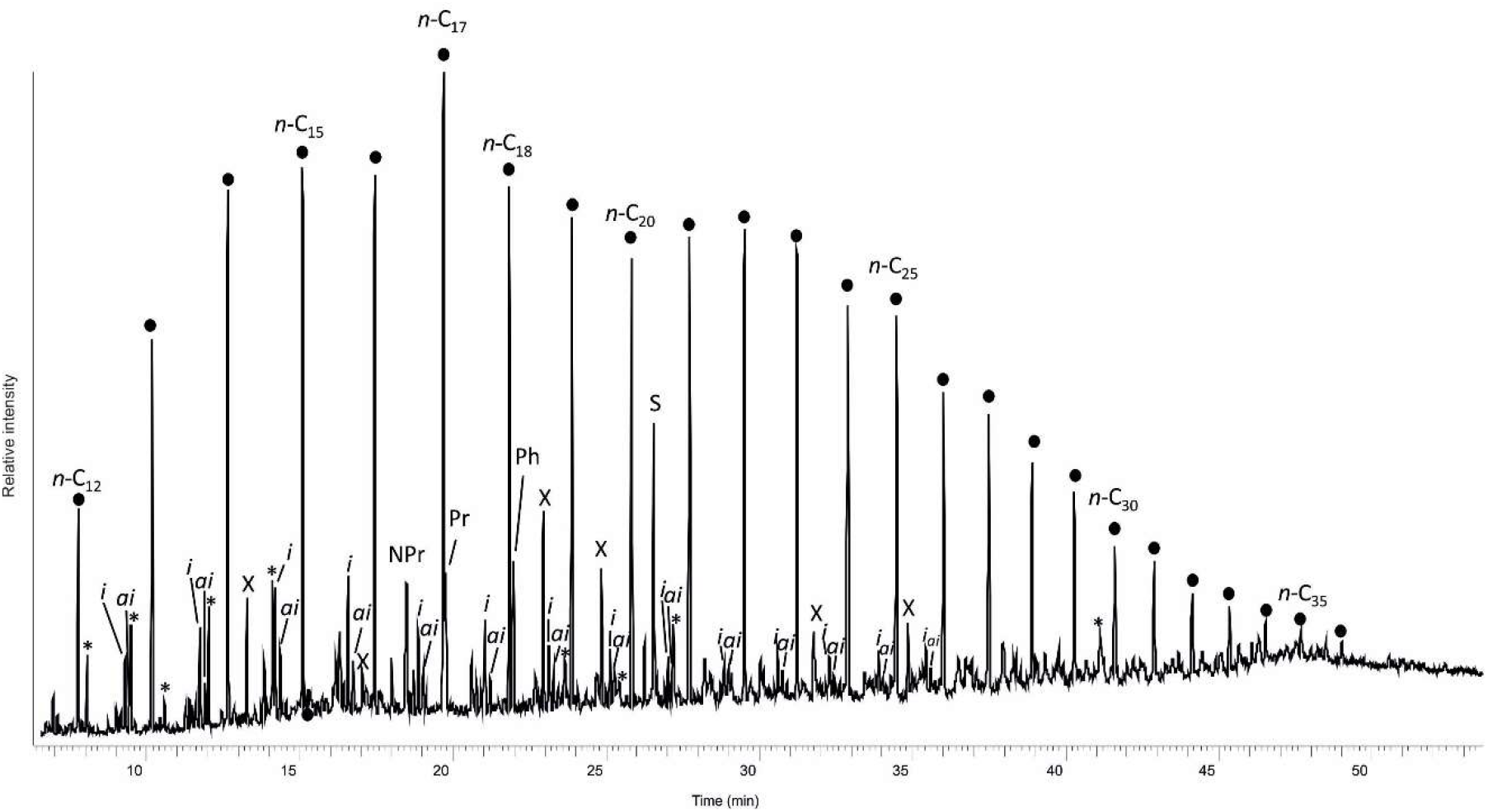
Total ion current chromatogram for a total organic extract (TOE) from a microbemetazoan build-up sampled at Mauer. Black circles = n-alkanes; *i* = iso-alkanes; *ai* = *anteiso-alkanes;* NPr = Norpristane; Pr = Pristane; Ph = Phytane; * = Unspecified acyclic isoprenoid; S = Elemental sulfur; X = Contaminant

### Carbon and oxygen stable isotopes

Carbon (δ^13^C_carb_) and oxygen (δ^18^O_carb_) stable isotope data for samples from Auerstedt and Libiąż cluster in three groups (Fig. 14). Group 1 comprises a bulk sample as well as 36 individual mineral phases and three *Placunopsis* bivalves in microbe-metazoan build-ups from Auerstedt. Group 2 includes 10 bulk samples from below and above the Auerstedt microbe-metazoan build-ups. Group 3 comprises 4 samples of individual mineral phases in Libiąż microbe-metazoan build-ups.

**Fig. 14.**
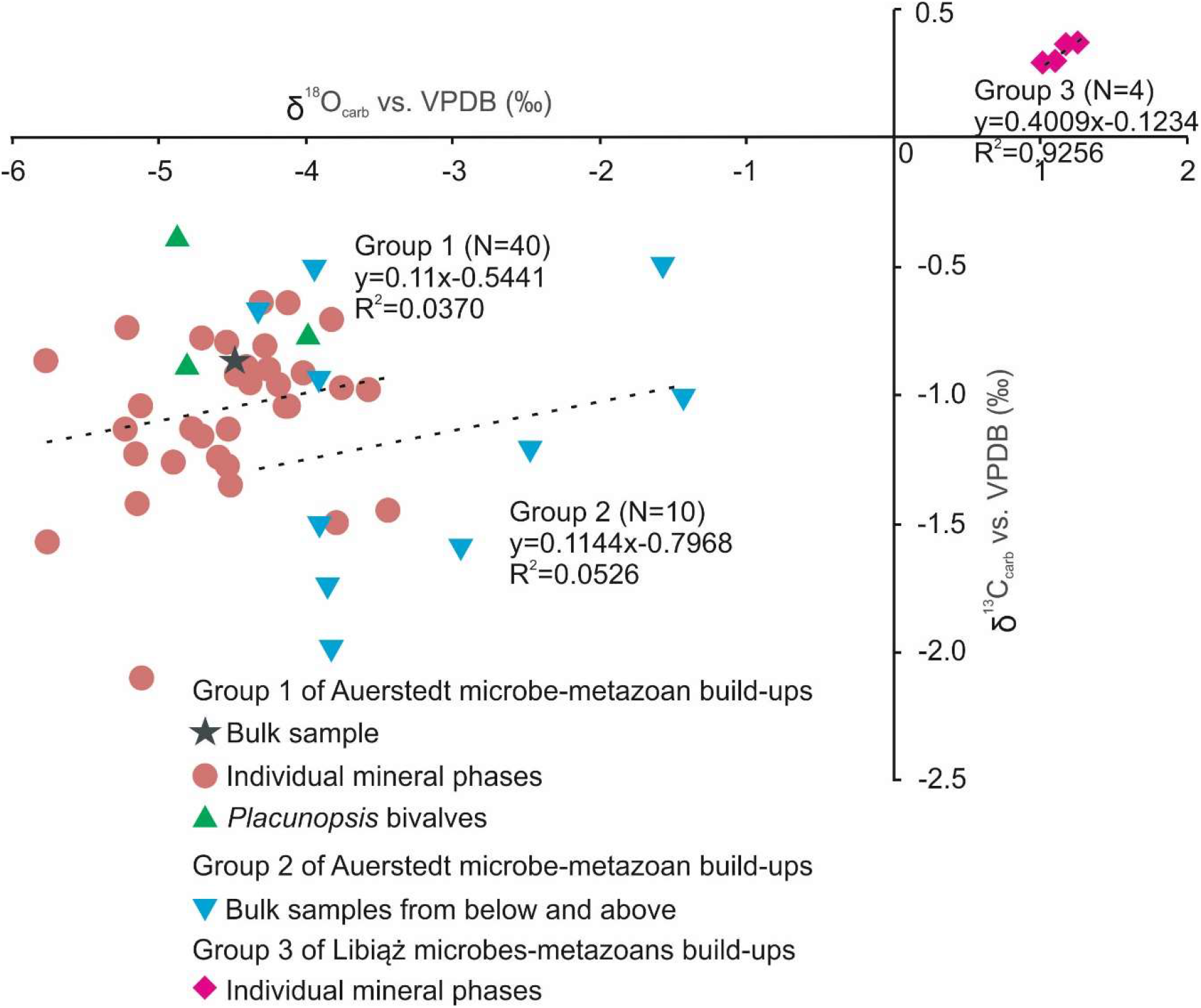
Carbon (δ^13^C_carb_) and oxygen (δ^18^O_carb_) stable isotope data for 54 samples from Auerstedt and Libiąż

Carbon (δ^13^C_carb_) and oxygen (δ^18^O_carb_) stable isotopes of Group 1 range from −2.1 to −0.4 and from −5.8 to −3.4 respectively (Fig. 14; Tab. 2). δ^13^C_carb_ values of Group 2 (between −2.0 and −0.5 are consistent with this record. δ^18^O_carb_ signatures of these samples, however, appear to be slightly less negative (between −4.3 and −1.4 (Fig. 14; Tab. 2). δ^13^C_carb_ and δ^18^O_carb_ values of Group 3 range from 0.3 to 0.4 and from 1.0 to 1.2 respectively (Fig. 14; Tab. 2), which is distinctly different to the data for samples from Auerstedt.

**Table 2.**
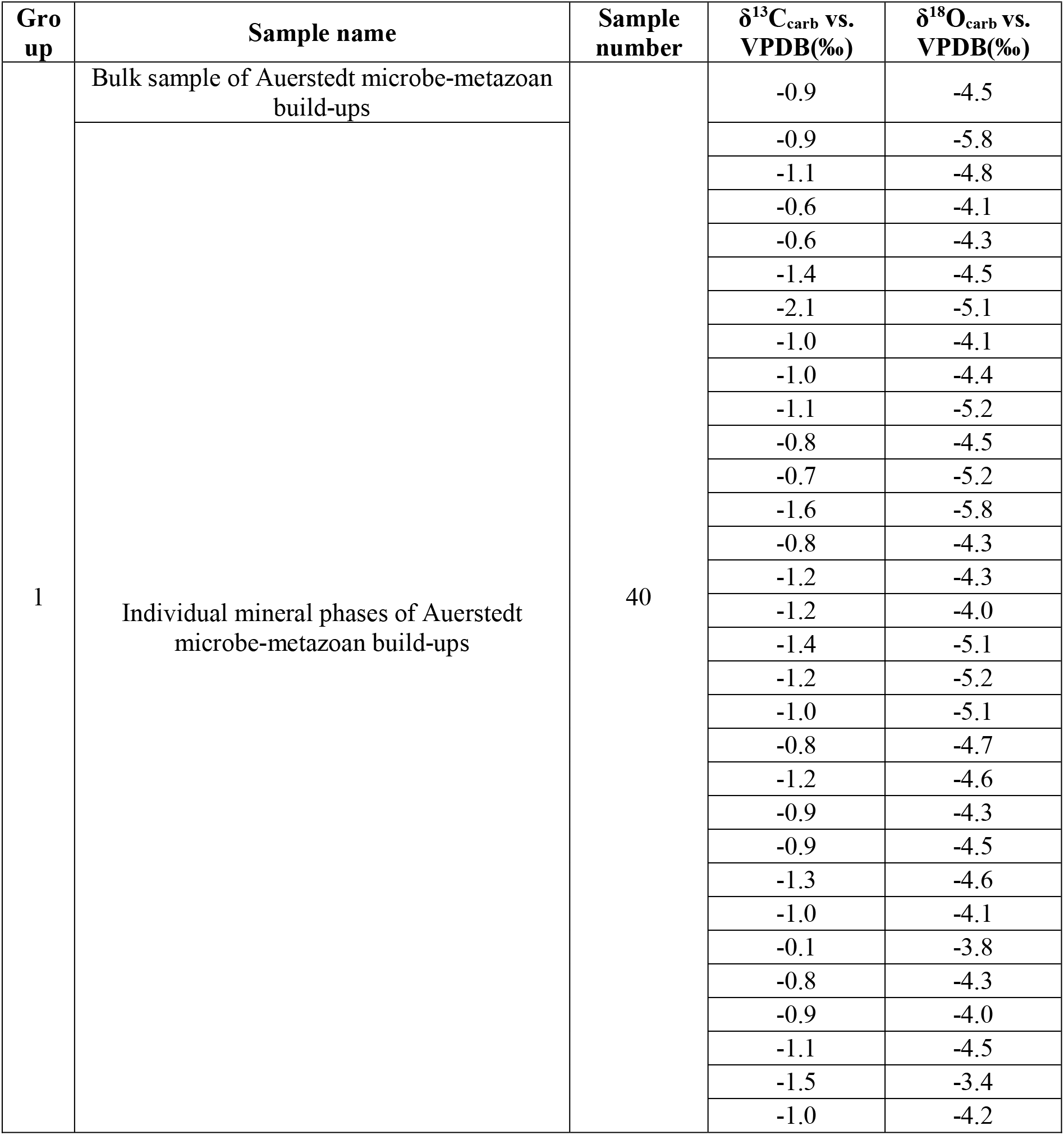
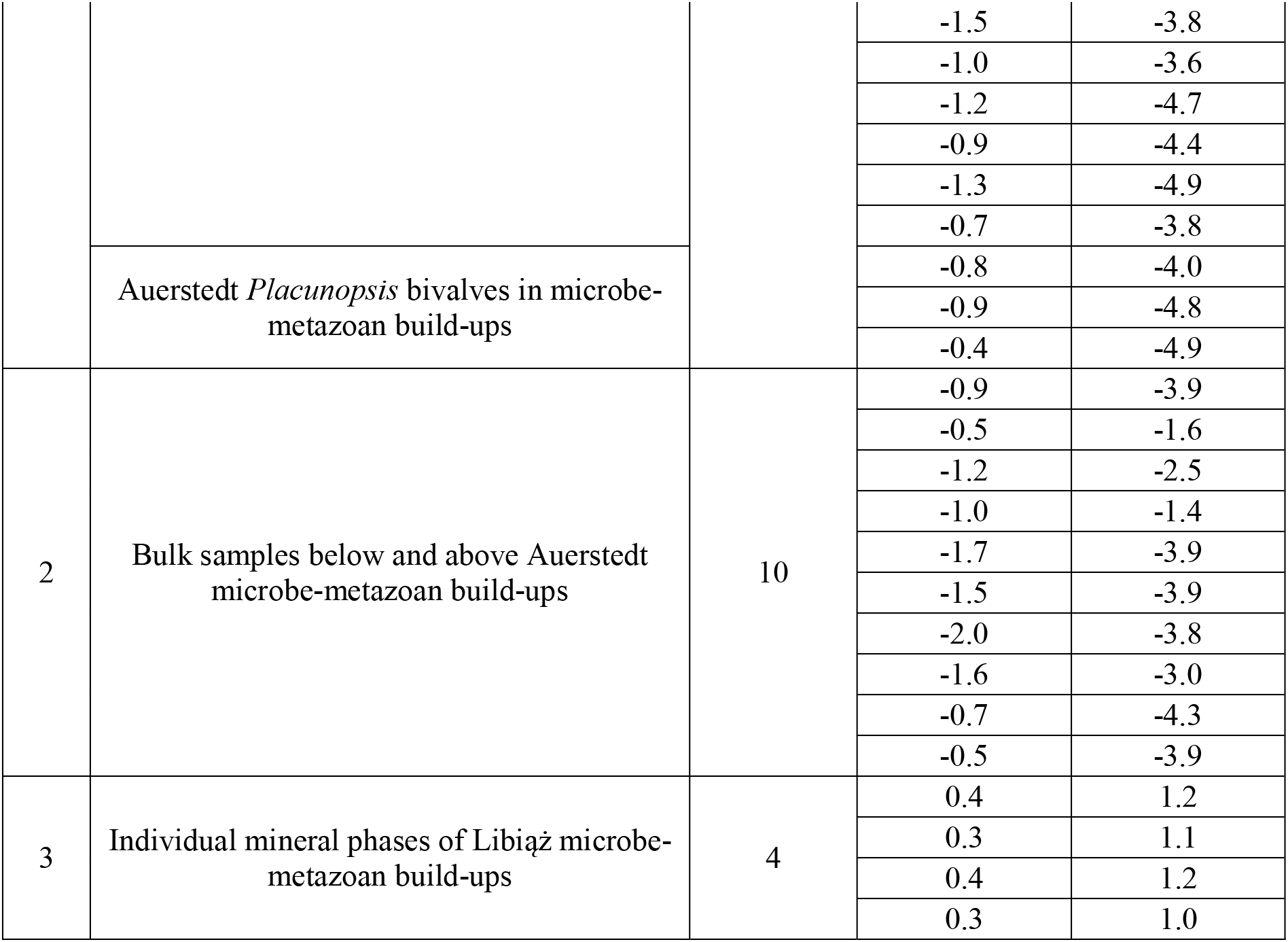
Carbon (δ^13^C_carb_) and oxygen (δ^18^O_carb_) stable isotope data for 54 samples from Auerstedt and Libiąż

δ^13^C_carb_ and δ^18^O_carb_ values of carbonates from Group 1 show a very low coefficient of determination (R^2^ = 0.0370; N = 40) (Fig. 14). The R^2^ value for stable isotope data from Group 2 is slightly higher but still low (R^2^ = 0.0526; N = 10). The low R^2^ values might indicate that these materials were barely affected by diagenetic influence. δ^13^C_carb_ and δ^18^O_carb_ values of Group 3, in contrast, exhibit a high coefficient of determination (R^2^ = 0.9256; N = 4) (Fig. 14), which may indicate a relatively strong diagenetic alteration (Bishop et al. 2014).

## Discussion

### Sedimentary environment of late Anisian microbe-metazoan build-ups

Paleogeographically, the Germanic Basin was a peripheral basin of the western Tethys Ocean during the Permian and Triassic periods (Götz & Feist-Burkhardt 2012; Ziegler 1990; Fig. 2). The presence of encrusting *Placunopsis* bivalves, *Spirobis-*like worm tubes in microbe-metazoan build-ups at Auerstedt (Fig. 6), as well as intraclasts, ostracods, gastropods, bivalves with borings and crustacean fecal pellets in the limestones atop (Figs. 3; 4c, d), indicate a shallow water environment. This interpretation is in good accordance with indications for elevated hydrodynamic energy regimes, such as disrupted laminated fabrics and randomly oriented bivalves in interspaces of microbe metazoan build-ups from Mauer, Seyweiler and Libiąż (Figs. 11c, d; 12b, e). Furthermore, δ^13^C_carb_ of about −2.1 to −0.4 in the analyzed samples are in the typical range of Triassic seawater (Veizer et al. 1999) (Fig. 14; Tab. 2). At the same time, the presence of calcite pseudomorphs after gypsum in the upper parts of microbe-metazoan buildups at Auerstedt (Fig. 10f, g) suggest elevated salinities. All these interpretations are in good accordance with an euryhaline, species-poor fauna in the Diemel Formation (Hagdorn & Simon 2005).

Taken together, the late Anisian microbe-metazoan build-ups were likely formed in a slightly evaporitic, subtidal shallow marine environment supporting previous work (Szulc, 1997; Krause & Weller 2000).

### Formation of late Anisian microbe-metazoan build-ups

The late Anisian microbe-metazoan build-ups in the Germanic Basin are characterized by diverse lamination types, including planar, wavy, domal and conical ones (Figs. 7; 10c–g; 11a–c; 12a–e). Layers that exhibit planar and wavy laminations were most likely mainly formed by microbial mats (Figs. 10c, e; 11a, b; 12a, e; Luo & Reitner 2016). Since the build-ups formed in very shallow marine environments and thus within the photic zone, cyanobacteria could have been the main microbes in the microbial mats. This is in good accordance with occurrences of pristane and phytane, which commonly derive from the phytol side chain of chlorophyll a and thus might reflect inputs by photoautotrophic organisms (e.g., Tissot & Welte 1984; Peters et al. 2005) (Fig. 13). Terminally branched alkanes (iso- and anteiso-alkanes), in contrast, possibly reflect contributions by sulfate-reducing bacteria (Kaneda 1991), which typically inhabit deeper layers in phototrophic microbial mats (Schneider et al. 2013). More detailed studies are needed to ensure that hydrocarbons in the investigated materials are indigenous and syngenetic to the host rock.

Layers that exhibit domal and conical laminations commonly show mesh-like fabrics and clotted to peloidal features (Figs. 10d, f, g; 11c, d; 12b–f). Similar features have been described from various Triassic build-ups across the Germanic Basin (Szulc 1997; Bachmann 2002; Luo & Reitner 2014, 2016). The interpretation of such features is controversial, ranging from an algal origin (Murray & Wright 1971) to a sponge origin (Szulc, 1997; Bachmann, 2002; Luo & Reitner 2014, 2016). The characteristics observed herein are remarkably similar to typical features of fossilized non-spicular demosponges (Luo & Reitner 2014, 2016). Particularly interesting in this regard are clotted to peloidal features, which are characteristic for automicrite formed *in situ* through the decay of microbe-rich sponge tissue (Reitner 1993; Reitner et al. 1995). Calcification of the sponge tissues during degradation may be attributed to sulfate reduction and ammonification, which potentially result in a locally increased carbonate alkalinity (Fritz 1958; Berner 1968; Reitner 1993; Reitner et al. 1995; Schumann-Kindel et al. 1997). Isolated pyrite crystals are also common in taphonomically mineralized sponge tissue (Reitner & Schumann-Kindel 1997). Notably, various modern demosponges (e.g., *Chondrosia reniformis*, *Petrosia ficiformis and Geodia barretti*) harbor abundant sulfate reducing bacteria (Reitner & Schumann-Kindel 1997S; Schumann-Kindel et al. 1997; Hoffmann et al. 2005). Based on Raman spectral images, three groups of dolomite can be distinguished in the meshlike fabrics and clotted to peloidal features (Fig. 9a, d–f). Dolomite of the first group is intimately associated with organic matter (Fig. 9d). The second group comprises clusters of anhedral dolomite crystals with varying orientations, as evidenced by small-sized variations of the intensity ratios of dolomite main bands in the Raman spectra (Fig. 9e). Dolomite of the third group is euhedral (Fig. 9f). These dolomite crystals exhibit preferred orientations, as suggested by consistent intensity ratios of the Raman bands. Dolomites of the first and second groups are suggested to attribute to protodolomite due to the intimate association with organic matter and the anhedral crystal shape. Protodolomite formation is perhaps linked to microbial sulfate reduction (Warthmann et al. 2000; Liu et al. 2020). It is therefore tempting to speculate that dolomites of the first and second groups reflect the activity of sulfate-reducing bacteria. Such an interpretation would agree with the presence of iso- and anteiso-alkanes (Kaneda 1991; see above).

Taken together, our findings reveal that the microbe-metazoan build-ups were formed by microbes (perhaps cyanobacteria and sulfate reducing bacteria) and metazoans (non-spicular demosponges), supporting findings from earlier studies (Szulc, 1997; Bachmann 2002; Luo & Reitner 2014, 2016). In addition to these organisms, *Placunopsis* bivalves and *Spirobis-*like worm tubes can contribute to the build-ups by inhabiting their upper surfaces once growth of microbes and non-spicular demosponges ceased.

### Paleoecological implications of late Anisian microbe-metazoan build-ups

The build-ups appear to be widespread in the Germanic Basin during Middle Triassic times, with occurrences in Baden-Wuerttemberg, Franconia, Thuringia, Brandenburg, Upper Silesia and the Holy Cross Mountains (Szulc 1997; Luo & Reitner 2014, 2016; Bachmann 2002). In addition, such build-ups are known from the direct aftermath of the Permian – Triassic Crisis in the Tethys realm (Friesenbichler et al. 2018; Heindel et al. 2018) and from Lower Triassic successions of the Northern Alps (Spötl 1988). The wide distribution of microbe-metazoan build-ups during Early to Middle Triassic times might imply a connection with the Permian – Triassic Crisis.

One possibility is that microbes and non-spicular demosponges filled the ecological niche cleared by the Permian – Triassic Crisis and benefited from a decreased ecological competition. A similar development is known from the aftermath of the Triassic – Jurassic mass extinction, when siliceous sponges reoccupied vacant shallow water environments from the deep (Delecat 2005; Delecat et al. 2011; Ritterbush et al. 2014). The investigated microbe-metazoan buildups, however, developed during late Anisian times (Middle Triassic), thus postdating the purported extinction event by about 8–9 Myr, when complex ecosystem was supposed to reemerge after the Permian – Triassic crisis (Chen & Benton 2012). Not like the outcrops in the Tethys realm (Friesenbichler et al. 2018; Heindel et al. 2018) as well as the Lower Buntsandstein “stromatolites” (Kalkowsky, 1908; Paul and Peryt, 2000; Paul et al. 2011) perhaps directly benefiting from the vacant niche after the crisis, the late Anisian build-ups maintained the advantage, indicating their resilience and adaptation to the ecosystem after the crisis.

Another, not necessarily contradictory possibility is that the environments were perhaps characterized by elevated salinities (Luo & Reitner 2016; this study). Microbial mats can develop under salinities of up to 170 (e.g. Schneider et al. 2013), but animals are commonly limited by high salinities (Bayly 1972). Notably, various sponges such as *Microciona prolifera* can survive in environments with salinities of up to 45 (e.g. Leamon & Fell 1990). Although it is unclear if non-spicular demosponges have the same tolerance, it seems plausible that these organisms profited from elevated salinities. Likewise, the alternate occurrences of microbial mats and non-spicular demosponges may also be controlled by changes in salinity (Luo & Reitner 2016).

Our observations support previous ideas that microbes and metazoans in the build-ups might have had a mutualistic relationship (Luo & Reitner 2016; Lee & Riding 2020). For all of these reasons, it is tempting to speculate that the investigated microbial-metazoan buildups reflect an ancient evolutionary and ecologic relationship. Unfortunately, it is challenging to explore further details on this issue, mostly because non-spicular demosponges have a relatively low fossilization potential due to the absence of spicules. The Anisian microbe-metazoan build-ups in the Germanic Basin, however, may help to track these organisms through geologic time and to clarify whether microbe-metazoan build-ups are more common than previously expected.

## Conclusion

The geobiology of late Anisian microbe-metazoan build-ups (upper Middle Muschelkalk) from four localities in the Germanic Basin (Auerstedt, Mauer, Seyweiler and Libiąż) were investigated. These build-ups consist of microbial mats, non-spicular demosponges, *Placunopsis* bivalves and/or *Spirobis-*like worm tubes. Therefore, these “stromatolites” should more correctly be referred to as microbe-metazoan build-ups. They are characterized by planar, wavy, domal and conical laminations. Notably, different lamination types reflect different organisms involved in their formation. Microbial mats likely played an important role in forming the planar and wavy laminations. Layers mainly built by non-spicular demosponges, in contrast, exhibit rather domal to conical laminations, mesh-like fabrics and clotted to peloidal features. The microbe-metazoan build-ups were obviously ecologically capable of coping with environmental change in the aftermath of the Permian – Triassic Crisis. It seems that the mutualistic relationship between microbes and non-spicular demosponges was the key to this success. They kind of maintained the advantage until the late Anisian. Another, not necessarily contradictory possibility is that they profited from elevated salinities. Due to the absence of spicules, non-spicular demosponges as well as the microbe-metazoan build-ups used to be overlooked in ancient rocks and should arouse increasing attention in the future.

## Acknowledgements

We thank Dr. Thomas Voigt for excellent support during fieldwork and providing papers, thank Dr. Gerhard Müller for providing silicified stromatolitic material from the Saarland area, and Dr. Hans Hagdorn for supporting us with additional rock material and valuable discussions. Johann Holdt is thanked for helpful field assistance. Axel Hackmann and Dennis Kohl are thanked for lab assistance. This study is financially supported by the China Scholarship Council.

